# Cryo-EM structures of Mycobacterium tuberculosis pyruvate carboxylase reveal allosteric activation and domain dynamics during catalysis

**DOI:** 10.1101/2025.11.11.687647

**Authors:** Aekagra Singh, Sandeep Singh, Uddipan Das

## Abstract

Pyruvate carboxylase (PC) is a biotin-dependent enzyme essential for anaplerotic flux into the tricarboxylic acid cycle and gluconeogenesis. Despite longstanding biochemical knowledge, the structural mechanisms governing its allosteric regulation and domain coordination remain poorly defined. Here, we present the cryo-EM structures of *Mycobacterium tuberculosis* PC (MtPC) captured in both apo and substrate-bound forms. In the absence of acetyl-CoA, MtPC forms a flexible and unstable tetramer that readily dissociates into subcomplexes. Upon binding of acetyl-CoA and the substrates, the enzyme transitions into a compact, ordered assembly, with acetyl-CoA acting as a molecular clamp that stabilizes inter-subunit interactions. Remarkably, we captured the mobile biotinylated domain simultaneously engaging each catalytic center within monomers, providing direct structural evidence for the two steps of the catalytic cycle. 3D variability analysis further revealed how this mobile domain translocates between the active sites during catalysis. Together, these findings demonstrate how acetyl-CoA stabilizes MtPC by reinforcing inter-subunit contacts and elucidate the conformational dynamics underlying domain coordination during catalysis. These findings provide new structural insights into the allosteric regulation of this essential metabolic enzyme.

## Introduction

Pyruvate carboxylase (PC, EC 6.4.1.1) is a highly conserved biotin-dependent enzyme that catalyzes the ATP-driven carboxylation of pyruvate to generate oxaloacetate^1^, a critical intermediate in the tricarboxylic acid (TCA) cycle in both bacteria and eukaryotes^2–5^. This reaction provides an essential anaplerotic pathway that replenishes TCA cycle intermediates and supports diverse metabolic processes, including gluconeogenesis, lipogenesis^3^, and neurotransmitter biosynthesis^6,7^. Structurally, PC is a single-chain, multi-domain enzyme of approximately 130 kDa that functions as a homotetrameric complex ^8–10^. Its activity relies on the interplay of three functional domains: biotin carboxylase (BC), carboxyltransferase (CT), biotin-carboxyl carrier protein (BCCP), and a central tetramerization (PT) domain that stabilizes the overall assembly.

Catalysis in PC proceeds through two sequential reactions, in which the BC domain first catalyzes the Mg-ATP-dependent carboxylation of biotin^11,12^, followed by the transfer of the activated carboxyl group to pyruvate by the CT domain, yielding oxaloacetate^13^. These reactions depend on the C-terminal BCCP domain, which carries a covalently attached biotin. The BCCP domain shuttles between the BC and CT active sites, spanning considerable distances between the distal domains within the tetrameric assembly^8,14–16^. Despite its central role in catalysis, the BCCP domain translocation pathway has not been directly observed yet. While large-scale domain rearrangements are known to underlie PC function, the dynamic coordination of these motions, and the cooperativity that emerges across the tetramer have remained elusive. Previous studies have captured only static snapshots of individual catalytic states^17^ or inferred transitions indirectly from kinetic data^18^, without resolving the continuous conformational changes that connect them.

PCs are allosterically regulated by acetyl-CoA, which binds at the junction of the PT and BC domains and stabilizes the enzyme in an active conformation^14–16,19^. The crystal structures of PC from Rhizobium etli (RePC) and Staphylococcus aureus (SaPC) have identified the acetyl-CoA binding site within the BC domain, providing key insights into the mechanism of allosteric regulation^8,14^. Cryo-EM studies of Homo sapiens PC (HsPC) have revealed conformational changes upon acetyl-CoA binding that stabilize the BC–BC dimer and enhance catalytic efficiency^20^. However, while HsPC structures demonstrate local stabilization, those of RePC and SaPC suggest that acetyl-CoA binding does not induce global conformational remodeling^15,21^. Although extensive biochemical studies have explored acetyl-CoA-mediated activation^14,22,23^, the precise molecular mechanism underlying its allosteric effect remains unclear. In particular, whether acetyl-CoA elicits localized structural changes or triggers broader reorganization across the tetramer remains unresolved, limiting our understanding of the regulatory principles and physiological roles of the PC.

Advancements in single-particle cryo-electron microscopy (cryo-EM) have transformed the study of macromolecular complexes at near-atomic resolution, offering unprecedented insights into their structural dynamics and functional flexibility^24,25^. Cryo-EM uniquely allows the visualization of heterogeneous conformational populations, capturing multiple functional states within a single dataset^26^. This is particularly advantageous for dynamic enzymes, such as PC, which undergo extensive domain movements during catalysis^27^. The advent of three-dimensional (3D) classification and 3D variability analysis (3DVA)^28^ further enhances this capacity by resolving subtle, continuous conformational transitions and revealing distinct catalytic intermediates^29^.

A detailed structural understanding of the dynamic transitions and regulatory mechanisms in PCs has remained elusive owing to the limitations in capturing transient and flexible states. Building on recent advances in cryo-EM and 3DVA, we applied these techniques to investigate the conformational landscape of *Mycobacterium tuberculosis* pyruvate carboxylase (MtPC) in both its apo and acetyl-CoA-bound forms. By combining ensemble-level visualization with focused structural refinement, we aimed to define the domain motions that underlie MtPC catalysis and examine how acetyl-CoA binding modulates tetramer stability and interdomain coordination. These structural insights may advance our understanding of allosteric regulation in dynamic multi-domain enzymes, particularly those requiring long-range domain coordination.

## Results

### Apo-MtPC adopts a flexible tetrameric architecture

To investigate the structural organization of MtPC, we cloned and overexpressed full-length MtPC in *E. coli* and purified it using affinity and size-exclusion chromatography. The chromatographic profile showed a predominant peak corresponding to the MtPC tetramer, along with a minor peak representing a dissociated species. SDS–PAGE analysis identified the latter as a degradation product, consistent with the mass of a CT–PT–BCCP fragment (Supplementary Fig. S1a), indicating that MtPC is prone to proteolytic cleavage and has a relatively short half-life. The dominant tetrameric fraction was used for cryo-EM analysis.

We first collected cryo-EM datasets of MtPC in the absence of acetyl-CoA, pyruvate, and ATP (hereafter referred to as the apo-MtPC). Single-particle analysis revealed two major particle populations: flexible intact apo-MtPC tetramers and stable CT-PT-BCCP dimers. The apo-MtPC tetramer was reconstructed at an overall resolution of 3.8-7.0 Å (Supplementary Fig. S2 and Supplementary Table 1) and adopts a square-shaped architecture measuring ∼160 × 167 Å (Fig. 1a). Similar to other PC, MtPC assembles as a homotetramer with BC–BC and CT–CT dimers arranged at opposite corners and PT domains directed toward the center (Fig.1b and Supplementary Fig. S1b–c). Each monomer comprises a BC domain with an internal B-subdomain insertion, a CT domain, and a split PT domain bridging these two. Owing to the limited resolution, the BCCP domain could not be modeled. However, the weak density near the CT domain suggests its approximate location (Fig. 1a). The CT and BC domains form the major inter-subunit interfaces. In contrast, the PT domains remain peripherally located and contribute minimally to tetramer stabilization, in contrast to RePC and SaPC, in which PT-PT interactions dominate tetramerization^8,14^. Notably, the B-subdomains of the BC regions were poorly resolved and adopted an open conformation, consistent with the destabilization of the ATP-binding site in the absence of nucleotides.

**Fig. 1:**
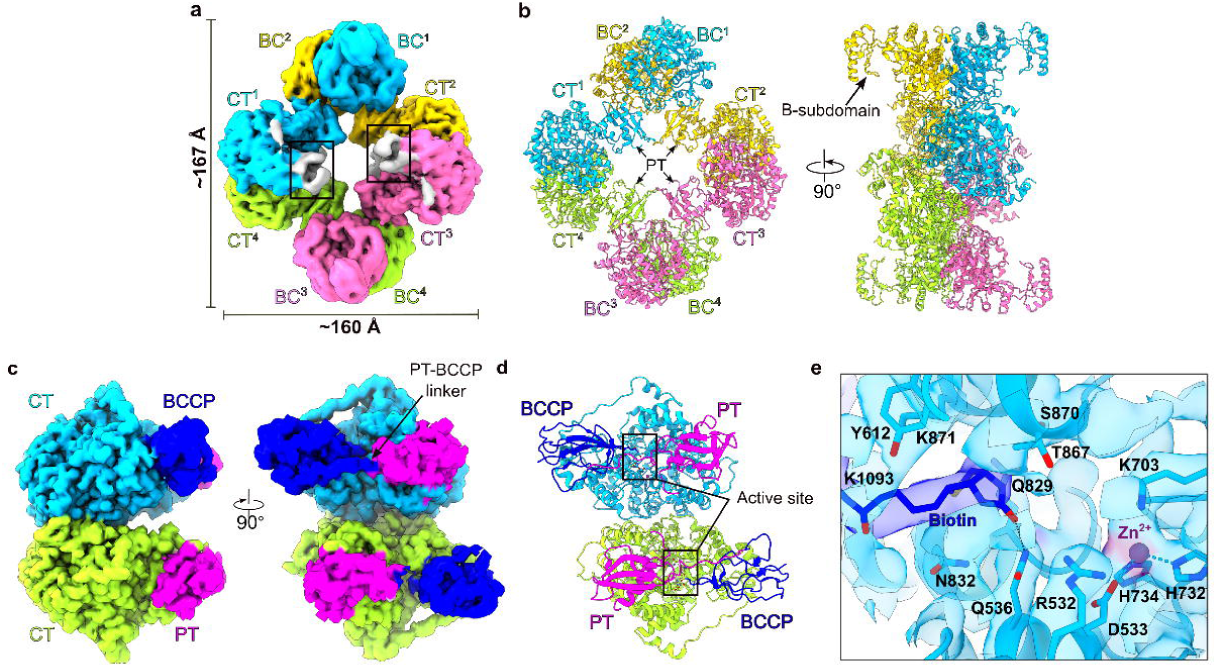
Structures of apo-MtPC and the CT–PT–BCCP subcomplex. **a** Cryo-EM map of apo-MtPC showing four monomers coloured deep sky blue, hot pink, yellow and light green, respectively, with the tetramer measuring approximately 160 × 167 Å. Insets highlight diffuse density corresponding to the BCCP domains near the CT regions. **b** Cartoon representation of the apo-MtPC tetramer showing the BC–BC and CT–CT interfaces at opposite corners, with the PT domains forming the central core. Orthogonal side views reveal the open configuration of the B-subdomain. **c** Cryo-EM map of the CT–PT–BCCP subcomplex. Colours follow the scheme used for the apo-MtPC tetramer, with the BCCP and PT domains shown in blue and magenta, respectively. Orthogonal views show the PT–BCCP linker and the antiparallel orientation of CT dimers. **d** Cartoon representation of the CT–PT–BCCP dimer showing the PT domain linked to the BCCP, which docks into the active-site pocket of its cognate CT domain. The PT and BCCP are coloured differently for clarity, though both belong to the same monomer. **e** Close-up view of the CT active site with bound BCCP. The covalently linked biotin (blue sticks) at K1093 forms a hydrogen bond with Q536. The Zn²⁺ ion at the catalytic site is coordinated by D533, H732 and H734, with K703 positioned toward the metal centre. CT-domain residues are shown within the cryo-EM density and coloured deep sky blue.

The second major particle class, corresponding to a CT–PT–BCCP dimer, was reconstructed at a resolution of 2.6-3.6 Å (Fig. 1c, Supplementary Fig. S2 and Supplementary Table 1). In this conformation, the CT domains form a dimer, with each BCCP domain connected to its corresponding PT domain through a flexible linker (Fig. 1d). The biotinylated K1093, located on the β-hairpin of the BCCP, inserts into the CT active site, while the BCCP body forms multiple stabilizing interactions, anchoring firmly to the CT lid (Supplementary Fig. S1d–e). The biotin forms a hydrogen bond with Q536 and lies in close proximity to Q829, N832, and T867, which together shape the CT active-site funnel (Fig. 1e). At the catalytic center, H732, H734, and D533 coordinate a divalent cation. Based on geometry and conservation, this ion was assigned as Zn²⁺, consistent with other PC structure^14^. No pyruvate density was detected, suggesting that this structure represents a degradation-stalled intermediate with biotin occupying the CT site in the absence of catalysis. The emergence of the dissociated CT–PT–BCCP dimer highlights the intrinsic instability of the apo-MtPC tetramer, which is highly flexible and prone to disassembly in the absence of substrates and cofactors.

### Substrate-bound MtPC adopts a compact conformation

To assess how substrate binding influences MtPC architecture, we determined its structure in the presence of acetyl-CoA, pyruvate, and ATP (hereafter referred to as substrate-bound MtPC). Initial refinement yielded a 2.9 Å map with C1 symmetry, which improved to 2.2–3.2 Å after applying D2 symmetry (Supplementary Fig. S3 and Supplementary Table 1). The resulting density was continuous for the BC, CT, and PT domains, although the B-subdomain of the BC domain remained disordered, reflecting intrinsic flexibility. Compared to the open, square-shaped apo-MtPC, the substrate-bound MtPC adopts a compact rhomboid architecture measuring ∼150 × 183 Å (Fig. 2a), which is stabilized by extensive inter-domain contacts involving the BC-BC, CT-CT, and PT-PT interfaces. Acetyl-CoA and ADP were resolved in the PT-CT junction and BC active sites, respectively, confirming that the enzyme was in a substrate-bound state during vitrification (Fig. 2b).

**Fig. 2:**
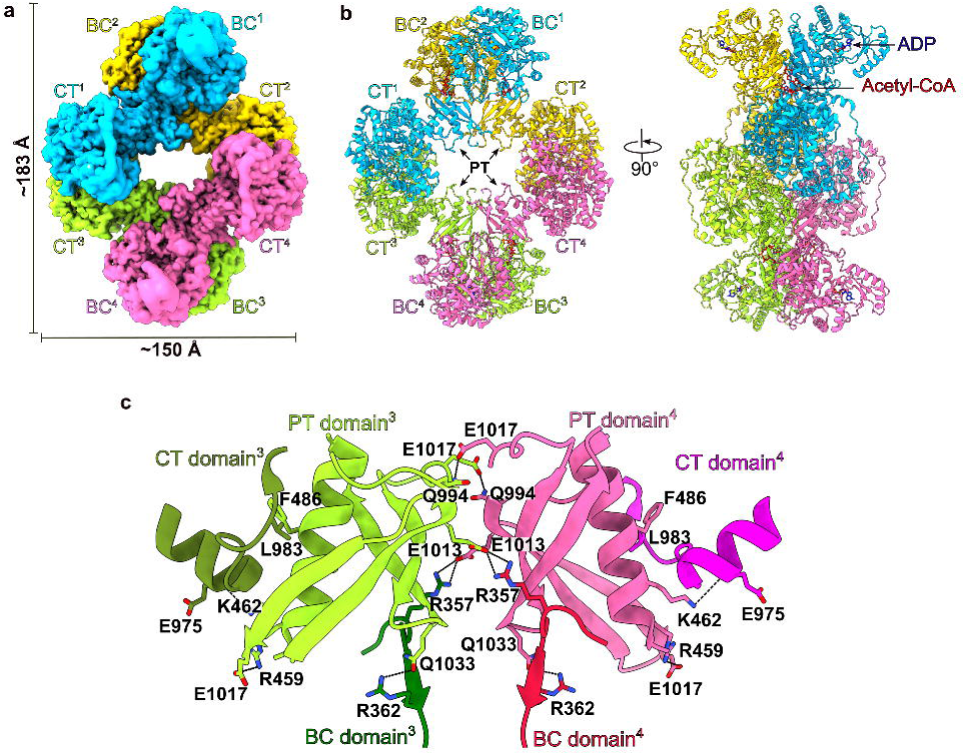
Cryo-EM structure of substrate-bound MtPC. **a** Cryo-EM map of MtPC, measuring approximately 150 × 183 Å, with four monomers coloured as in the apo-MtPC structure. **b** Cartoon representation of the MtPC tetramer showing a compact rhomboid architecture with closely positioned PT–PT domains. Orthogonal side views highlight acetyl-CoA (red sticks) and ADP (blue sticks) bound at the BC–CT hinge region and within the B-subdomain, respectively. **c** Interactions between adjacent monomers mediated by their PT domains, coloured light green and hot pink, respectively. The associated BC (dark green and red) and CT domains (olive green and magenta) are shown in distinct colours for clarity. Hydrogen bonds (≤ 3.5 Å) are indicated by black dashed lines. Domain labels correspond to their respective colours, and superscripts ³ and ⁴ denote adjacent monomers.

A detailed analysis revealed that the BC domain forms stabilizing contacts with both termini of the PT domain (453-478, 996-1045). A loop in the BC domain (residues 359–377) engages the PT tail via a hydrogen bond between R362 and Q1033 and a bidentate salt bridge between R357 and E1013 that extends across the adjacent monomer (Fig. 2c). The PT domain also interacts extensively with the CT domain: the N-terminal PT helix aligns along the CT surface, forming a hinge, whereas R459 hydrogen bonds with E1003, stabilizing the acetyl-CoA binding cleft. Additional contacts, including L983–F466 and K462–E975, further anchored the PT to the CT interface. At the tetramer core, the C-terminal segment of PT extends as an antiparallel four-stranded β-sheet, with interchain hydrogen bonds between Q994 and E1017 forming a reciprocal seam, thereby reinforcing the PT–PT interface as a central stabilizing interaction (Fig. 2c).

Superimposition of the apo and substrate-bound MtPC structures using one chain as a reference revealed domain-specific conformational shifts (Fig. 3a and Supplementary Movie 1). In the apo-MtPC, the CT lid of monomer 1 tilts by approximately 23° toward the CT funnel, accompanied by a ∼26° rotation of the BC domain, while the PT domain remains largely stationary (Fig. 3b). The adjacent monomer-2, linked via the BC–BC interface, is rotated by ∼15°, producing the square-shaped arrangement observed in apo-MtPC (Fig. 3c). Conversely, nucleotide binding in the substrate-bound MtPC induces a ∼44° closure of the B-subdomain (Fig. 3d). The PT domains also undergo significant rearrangement in apo-MtPC, where the inter-PT distance increases by ∼9 Å (Fig. 3e) as the adjacent PT domain rotates ∼10° away from the tetramer plane (Fig. 3f), disrupting the central PT–PT seam. Despite these rearrangements, the apo structure retained the tetrameric assembly, indicating that tetramer formation can occur independently of PT-PT interactions.

**Fig. 3:**
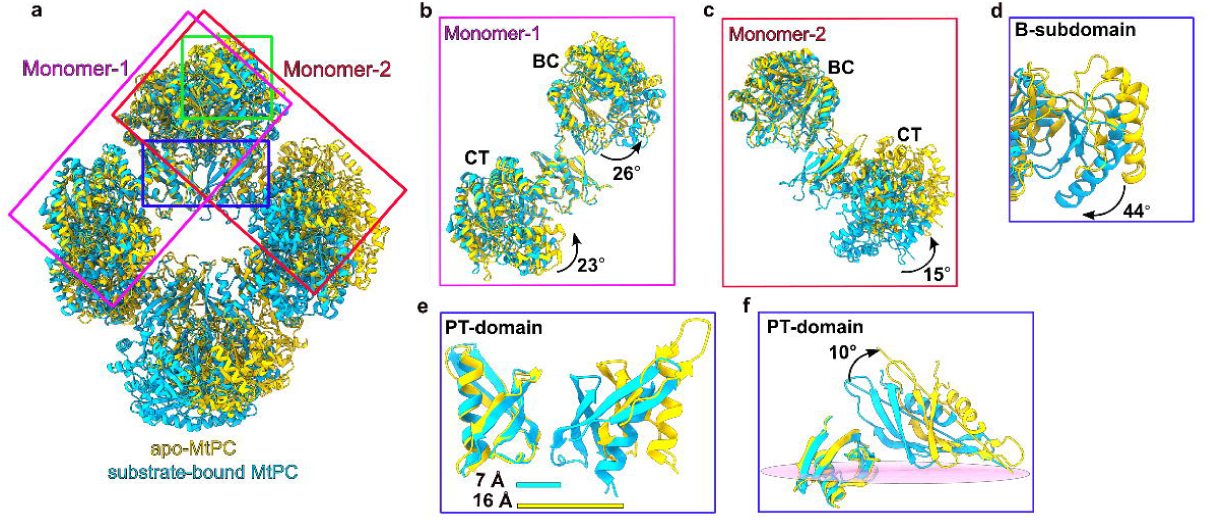
Overlay of apo and substrate-bound MtPC. **a** Overlay of apo-MtPC (yellow) and substrate-bound MtPC (deep sky blue) cartoon models. The front monomers are outlined by a magenta rectangle, monomer 2 by a red rectangle, and the PT–PT interface and B-subdomain by blue rectangles. **b** Overlay of monomer 1 showing the CT lid in apo-MtPC tilted by ∼23° toward the CT funnel, while the BC domain rotates by ∼26°. **c** Overlay of monomer 2 showing a ∼15° tilt of the CT domain relative to the BC domain in apo-MtPC. **d** The B-subdomain in MtPC exhibits a ∼44° closure relative to apo-MtPC. **e** Overlay of two adjacent monomers showing PT domains separated by ∼16 Å in apo-MtPC compared with ∼7 Å in substrate-bound MtPC. **f** Relative rotation of the apo-MtPC PT domain by ∼10° compared with substrate-bound MtPC, resulting in an increased PT–PT interface distance. See also Supplementary Movie 1 for the full conformational transitions.

### Acetyl-CoA binding stabilizes the tetrameric architecture

To assess the catalytic effect of acetyl-CoA, MtPC activity was measured in the presence and absence of the allosteric activator under steady-state conditions. In the absence of acetyl-CoA, MtPC displayed a specific activity of 3258 min⁻¹, which increased to 4428 min⁻¹ upon addition of the cofactor, corresponding to a ∼1.4-fold enhancement in activity (Fig. 4a). Michaelis-Menten analysis further revealed that acetyl-CoA increases the *k*_cat_ from 64.1 s⁻¹ to 77.7 s⁻¹, while lowering the apparent *K*_m_ for pyruvate from 2.31 mM to 1.49 mM, indicating improved substrate affinity (Fig. 4b). Together, these results demonstrate that acetyl-CoA enhances MtPC catalysis both by stabilizing the active tetramer and by promoting efficient substrate channeling between the BC and CT active sites.

**Fig. 4:**
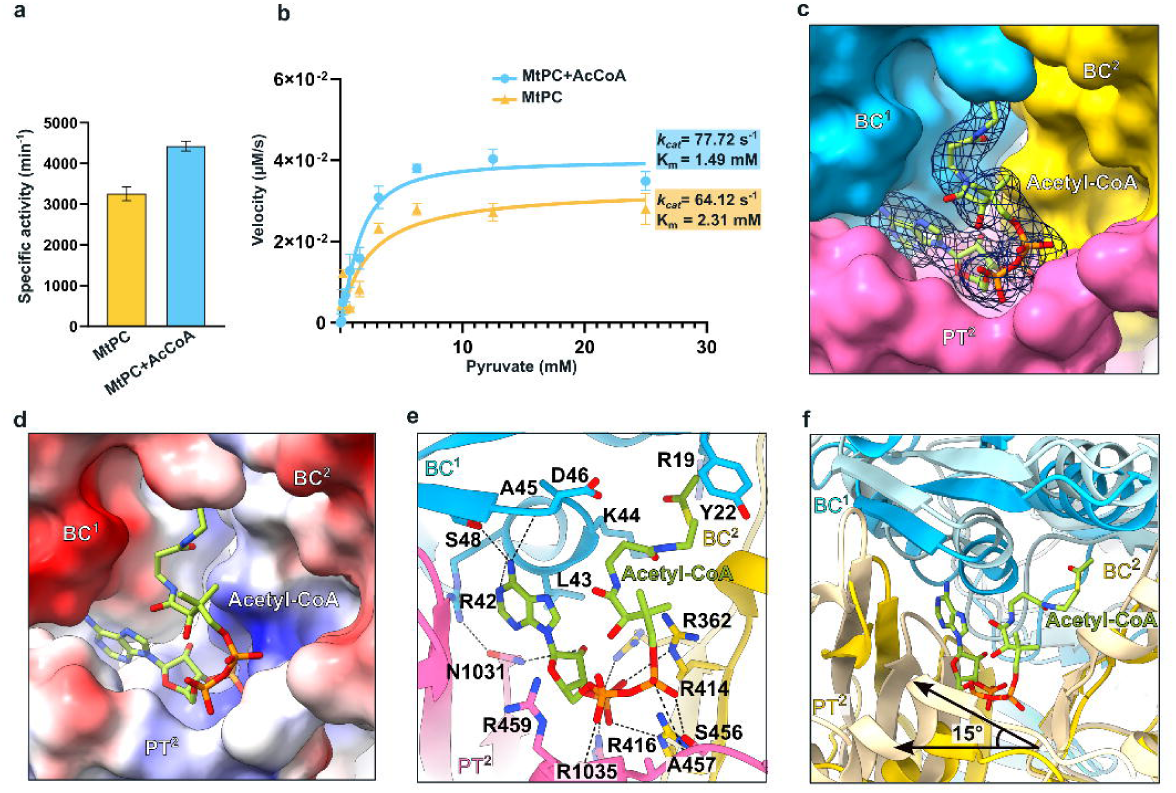
Acetyl-CoA binding site and allosteric regulation of MtPC. **a** Specific activity of MtPC in the presence and absence of Acetyl-CoA. Bars represent mean values from biological triplicates, with error bars showing standard error of the mean (SEM). Statistical significance was determined using an unpaired two-tailed t-test with Welch’s correction (P = 0.0068). **b** Enzyme kinetics of MtPC measured using the PC–MDH coupled assay. Data were collected from biological triplicates and fitted to an allosteric model for the Acetyl-CoA (+) condition and a Michaelis–Menten model for the Acetyl-CoA (–) condition. **c** Surface representation of the acetyl-CoA-binding cavity formed at the interface between adjacent monomers 1 and 2. Monomer 1 is coloured deep sky blue and monomer 2 yellow, with the PT domain of monomer 2 highlighted in hot pink. Acetyl-CoA is shown as green sticks, with the corresponding cryo-EM density displayed as a dark-blue mesh. **d** Electrostatic surface potential of the acetyl-CoA-binding pocket, coloured from –10 kT e⁻¹ (red) to +10 kT e⁻¹ (blue). **e** Close-up view of residues from the BC domains of adjacent monomers and the PT domain involved in acetyl-CoA binding. Hydrogen bonds (≤ 3.5 Å) are indicated by black dashed lines. **f** Overlay of the acetyl-CoA-binding site in apo-MtPC (BC¹ cyan; PT and BC² wheat) and substrate-bound MtPC (BC¹ deep sky blue; PT and BC² yellow), with domain shifts indicated by angle. See also Supplementary Movie 2 for the conformational differences. Graph plotting, curve fitting, kinetic parameter determination (*K*_m_ and *k*_cat_), and statistical analyses were performed using GraphPad Prism 10.6.1.

Our structural analysis revealed that acetyl-CoA binds at the junction of the BC dimer and PT domain, bridging adjacent BC monomers and extending into the PT groove (Fig. 4c). Electrostatic surface mapping revealed that one BC domain contributes a negatively charged patch, whereas the adjacent BC and PT domains form positively charged grooves that cradle the acetyl-CoA molecule (Fig. 4d). This binding site corresponds to the conserved allosteric pocket previously described in other PC studies^20^.

In substrate-bound MtPC, the adenine moiety of acetyl-CoA is anchored to the BC domain via hydrogen bonds with the backbone of A45 and the side chain of S48, whereas R42 stabilizes the nucleotide pocket through an interaction with N1031 of the PT domain. The 3′-phosphate is enclosed by an arginine-rich cage composed of R362, R414, R416, and R1035 from the adjacent BC monomer, mirroring the conserved residues found in SaPC and HsPC. The pantetheine arm follows the PT groove, interacting with S456, A457, and R459 before docking into a hydrophobic pocket in the neighboring BC monomer, where D46, K44, Y22, L43, and R19 form stabilizing contacts (Fig. 4e). Together, these three anchoring elements-nucleotide coordination, phosphate encapsulation by the arginine cage, and pantetheine docking, form an interdomain bridge between the BC and PT domains, thereby reinforcing the PT–PT seam at the tetramer core.

Structural overlay of the acetyl-CoA binding pocket of apo and substrate-bound MtPC revealed marked conformational differences. In the absence of acetyl-CoA, the groove at the BC-PT-BC junction was open, with the BC domains misaligned relative to the PT domains (Fig. 4f, Supplementary Movie 2). Acetyl-CoA binding pulls the BC and PT domains of one monomer closer to the BC domain of the adjacent monomer, reinforcing the interdomain groove and promoting a compact assembly. Acting as a molecular clamp, acetyl-CoA stabilizes inter-subunit alignment and locks the tetramer into a catalytically competent rhomboid configuration. These findings establish acetyl-CoA as a key structural determinant of MtPC tetramer stability and architectural integrity.

### BCCP domains can simultaneously engage their BC and CT active sites

The dynamic translocation of the BCCP domain is central to the PC catalytic cycle but has historically eluded direct structural visualization because of its flexible loop connecting to the CT domain. In our initial C1-symmetry reconstruction of the substrate-bound MtPC, we observed diffuse density bridging the BC and CT active sites which is indicative of conformational heterogeneity arising from BCCP mobility. To better resolve this region, we applied symmetry expansion, followed by 3D classification (Supplementary Fig. S4a). This approach partitioned the dataset into ten classes, of which ∼65% exhibited resolved BCCP density, while ∼35% lacked detectable BCCP signals, likely reflecting rapid conformational exchange (Supplementary Fig. S4c–d). Among the BCCP-resolved classes, the domains were consistently localized to two positions, designated site 1 and site 2, corresponding to alternate engagement with the BC and CT active sites within the tetramer (Fig. 5a). Class 4 (∼17%) showed BCCP from monomers 1 and 2 docked at site 1 on the front face and site 2 on the back face. Class 6 (∼16%) displayed exclusive occupancy at site 2 with undetectable density at the back face, whereas classes 8 and 9 (∼17% and ∼15%, respectively) captured at site 1 and site 2, respectively, across both tetramer faces.

**Fig. 5:**
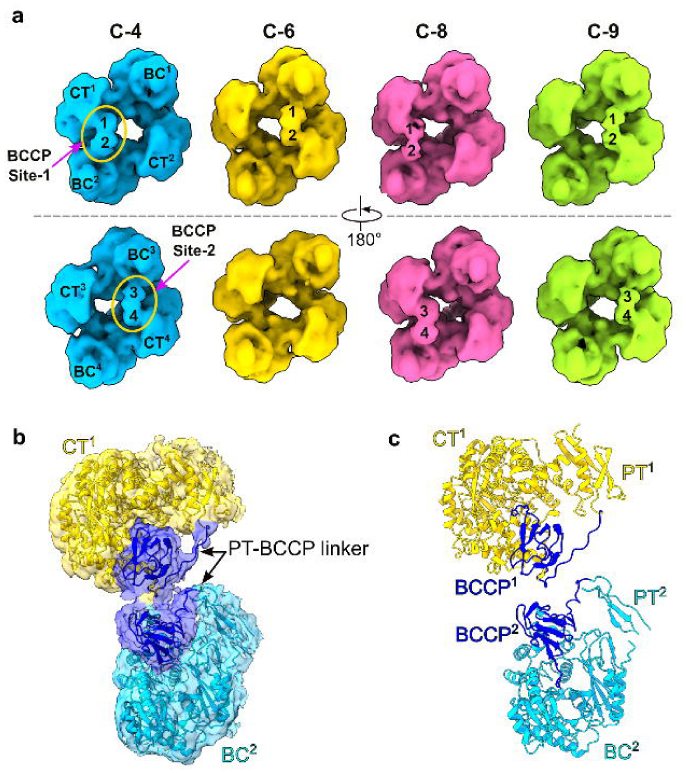
3D classification of MtPC. **a** Cryo-EM maps of substrate-bound MtPC after 3D classification showing four representative classes (C4, C6, C8 and C9) coloured deep sky blue, yellow, hot pink and light green, respectively. Each class displays a distinct arrangement of BCCP domains; superscripts denote monomer identity. Upper panels show front views and lower panels show back views rotated 180°. **b** Cryo-EM map of the focused class with fitted atomic model, showing adjacent monomer densities with BCCP domains (blue) engaged in the BC (deep sky blue) and CT (yellow) active sites, with continuous density extending from the parent monomer toward the BCCP domain. **c** Cartoon representation of the BC–BCCP and CT–BCCP fitted models illustrating inter-domain connections and relative positioning of the catalytic sites..

To further improve the resolvability and enable accurate modeling of the BCCP within these sites, we applied a focused mask around only site 1 (as sites 1 and 2 are structurally equivalent) and subtracted the remaining tetramer signal before a second round of 3D classification. This method isolated ∼21% of the particles with a clear BCCP density (Supplementary Fig. S4e). Local refinement of this subset yielded a 2.9 Å map (Fig. 5b, Supplementary Fig. S4b and Supplementary Table 1), enabling the reliable modeling of two BCCP domains simultaneously docked at the BC and CT sites of their respective parent monomers (Fig. 5c). This structural state, which features dual intra-monomer BCCP engagement, has not been reported in previous PC structures. Notably, the refined density unambiguously resolved the loop linking each BCCP to its PT domain, confirming that BCCP translocation in MtPC can occur within the same monomer.

### Structure of BCCP engaged at BC and CT active sites

The refined BC–BCCP/CT–BCCP model offers detailed insights into the active centers of MtPC. In the BC domain, a loop from the PT domain of the parent monomer extended toward the BCCP, positioning biotinylated K1093 within the nucleotide-binding pocket (Fig. 6a). The BCCP core is flexibly tethered, forming only transient contacts with the BC domain, which is consistent with its dynamic nature. Accordingly, biotin density was only visible at lower map thresholds (Supplementary Fig. 5a). In contrast, the BC domain itself adopts a well-ordered ATP-grasp fold, with the B-subdomain captured in a closed conformation, acting as a molecular lid that regulates nucleotide access. The surrounding residues were well resolved, enabling confident modeling of the nucleotide-binding pocket.

**Fig. 6:**
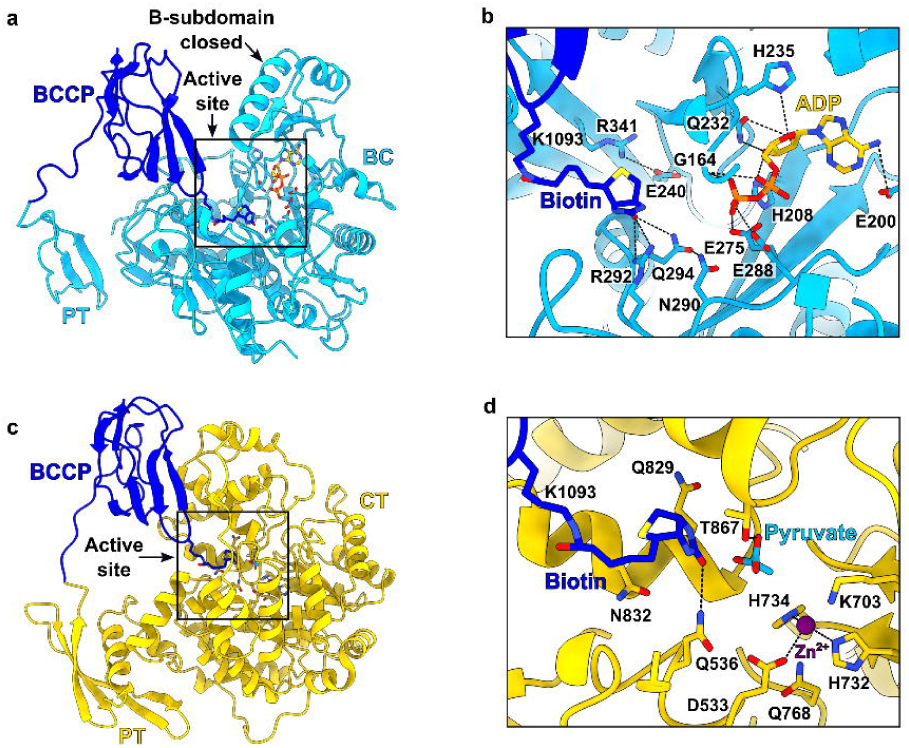
Structural details of BC–BCCP and CT–BCCP interactions. **a** Cartoon representation showing interaction of the BCCP (blue) with the BC domain (deep sky blue). The BCCP belongs to the same monomer and is connected via the PT–BCCP linker. Biotin (blue) and ADP (yellow) are positioned within the active site, which adopts a closed B-subdomain conformation. The boxed region indicates the active site. **b** Close-up of the BC active site showing biotin juxtaposed near the ADP-binding pocket, with interacting residues within 3.6 Å shown. **c** Interaction of the BCCP with the CT domain (yellow). The boxed region highlights the active site with biotin covalently attached to K1093 of the BCCP. **d** Close-up of the CT active site showing the Zn²⁺ (purple) coordination centre and biotin interacting with Q536, while pyruvate (cyan) interacts with T867.

In this conformation, biotin forms a hydrogen bond with R292 and is further oriented by Q294 and N290 to align the reactive N1 atom for catalysis (Fig. 6b). ADP is stably bound to the BC site: the adenine ring interacts with E200; the ribose contacts H208, H235, and Q232; and the polyphosphate coordinates with E275, E288, and the main chain of G164 in the T-loop. Interestingly, biotin lies ∼6.1 Å from the β-phosphate of ADP, consistent with the post-hydrolysis state. This is further supported by the presence of a salt bridge between R341 and E240, a gatekeeping interaction proposed to block carboxybiotin from re-entering the BC site and preventing premature decarboxylation^30^. Although limited local resolution precluded the definitive assignment of carboxybiotin, the structural features support a post-carboxylation intermediate.

Our locally refined map also resolved a BCCP domain engaged with its monomer at the CT-active site (Fig. 6c). The biotinylated β-hairpin of BCCP is inserted into the CT domain, positioning K1093 within the active site for catalysis. This insertion is stabilized by hydrophobic interactions between the BCCP core (residues 1095-1101) and a loop in the CT lid (residues 896–901), which is part of the C-terminal extension of the TIM-barrel fold. The CT lid adopts a closed conformation, analogous to that observed in the isolated CT-PT-BCCP dimer (Supplementary Fig. 1d-e), supporting a regulatory role for the lid in BCCP engagement. The loop connecting BCCP to the PT domain was well resolved, confirming intra-monomeric translocation (Fig. 5c).

Within the CT active site, biotin is coordinated by residues Q829, N832, and T867. Although its density is weaker than that of the surrounding residues, as observed in previous PC structures^8^, (Supplementary Fig. 5b), it remained clearly traceable. The catalytic site, embedded within the TIM-barrel fold, was well resolved, enabling accurate modeling of key active-site residues. A pyruvate molecule was positioned adjacent to biotin, interacting with T867 near the Zn²⁺ ion, which is coordinated by H732, H734, D533, and K703 (Fig. 6d). This configuration closely mirrors that observed in RePC and captures the CT domain in a catalytically poised state, ready for carboxyl transfer^14^.

### 3D variability analysis reveals coordinated BCCP motion across the catalytic cycle

To capture the continuous conformational landscape of substrate-bound MtPC during catalysis, we performed 3DVA on symmetry-expanded C1 particles using a 3 Å filter resolution. The analysis revealed a continuous distribution of conformational states along the principal components of motion, indicating that substrate-bound MtPC samples a dynamic ensemble rather than adopting discrete structural intermediates (Fig. 7a).

**Fig. 7:**
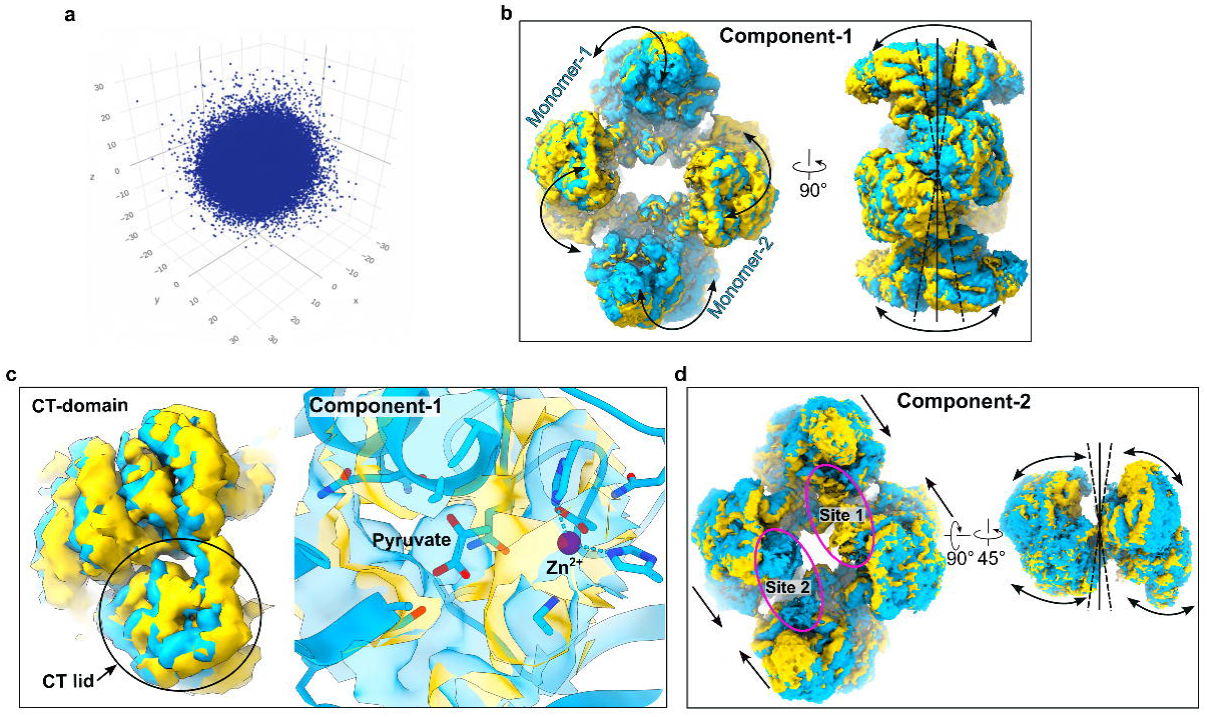
3D variability analysis of MtPC. **a** Scatter plots of 3DVA reaction coordinates of individual MtPC particles showing the range of structural variability along sequential component pairs. **b** Two extreme trajectories from 3DVA Component 1 (first frame, deep deep sky blue; tenth frame, yellow) reveal a see-saw motion of the top and bottom BC dimers, accompanied by CT dimers moving alternately toward and away from the tetramer centre. Orthogonal views illustrate the relative domain movements, with arrows indicating the direction of motion. **c** Relative shift of the CT lid between the two extreme trajectories, with corresponding cryo-EM densities showing pyruvate in the first trajectory (deep sky blue) and its disappearance in the last (yellow). **d** Component 2 captures bending motions between monomers 1 and 2, oscillating toward and away from each other, accompanied by the appearance of BCCP domain densities (marked by magenta ellipses). Guide lines from the map centre serve as visual reference points. See also Supplementary Movie 3 and 4 for the full conformational transitions.

The first principal component (or eigenvector) represented the dominant trajectory, characterized by a see-saw motion of the BC dimers pivoting around the PT domain hinges (Fig. 7b; Supplementary Movie 3). This movement was coupled with the reciprocal bending of the CT dimers, such that the front tilting of the BC dimers toward the tetramer center coincided with the outward displacement of the CT dimers. This component also revealed the loss of detectable pyruvate density with the inward tilting of the CT lid, consistent with oxaloacetate release and resetting of the catalytic cycle (Fig. 7c). The second component captured a lateral bending motion between the front-facing monomers and resolved the weak BCCP density at sites 1 and 2 (Fig. 7d; Supplementary Movie 4). Interestingly, this mode uncovered a coordinated pattern of BCCP movement that mirrored the transitions identified in the 3D classification. As the front-facing BCCP domains alternated between sites 1 and 2, the back-facing BCCPs disengaged and transitioned toward the diagonally opposite active sites. This reciprocal motion demonstrates that all eight active sites within the tetramer are functionally interconnected, offering structural support for a highly coordinated BCCP translocation cycle in MtPC.

Although the high-resolution consensus map lacked sufficient BCCP density for reliable modeling, 3DVA resolved its transient engagement states, providing mechanistic insight into the domain’s intrinsic flexibility. To capture the full trajectory of BCCP motion, a second 3DVA was performed with a 12 Å resolution cutoff, yielding a lower-resolution map that clearly revealed the BCCP density and its continuous connection to the parent PT domain. In the first component, each BCCP was docked at site 1 on both faces in one conformation and at site 2 on both faces in another. The second component revealed a reciprocal arrangement, with BCCP engaging site 1 on the front face and site 2 on the back face (Supplementary Fig. 6a, 6b; Supplementary Movie 5). These observations are consistent with our 3D classification results but, importantly, 3DVA provided a dynamic view of the continuous transition between these states, capturing the path of BCCP translocation between the two catalytic centers. Collectively, these analyses delineate the complete trajectory of BCCP motion, revealing a coordinated and alternating translocation mechanism that underpins the MtPC catalytic cycle.

## Discussion

PC is a key metabolic enzyme that catalyzes a multi-step reaction involving long-range domain translocation; however, capturing its dynamic conformational landscape remains challenging. Previous studies, including cryo-EM analyses of LiPC, revealed distinct catalytic states^17^, whereas the structures of HsPC underscored the allosteric role of acetyl-CoA in stabilizing the BC domain and promoting tetramer assembly^20^. However, these snapshots do not resolve the continuum of structural transitions underlying catalysis. Here, by combining high-resolution cryo-EM with 3DVA, we present an integrated structural framework for MtPC, delineating its transition from a flexible apo assembly to a compact, catalytically competent tetramer. Interestingly, we resolved the sequential engagement of the BCCP domain with both active sites of the same monomer, providing direct visualization of its coordinated translocation and uncovering key intermediates that define the dynamic reaction cycle of the enzyme.

In the apo state, MtPC adopts a flexible, square-shaped tetrameric structure in which the PT domains are displaced from the core, leading to weakened inter-subunit contacts. This contrasts with the canonical quaternary architectures of RePC^14^ and SaPC^8^, where the PT domains contribute to critical stabilizing interactions. The propensity of the apo MtPC tetramer to dissociate into CT-PT-BCCP dimers highlights its inherent instability in the absence of cofactor. This behavior parallels observations in apo HsPC, where acetyl-CoA depletion results in poorly resolved BC domains^20^. Collectively, these findings support a conserved role for acetyl-CoA in maintaining the structural integrity of the PC tetramer.

Upon acetyl-CoA binding, MtPC undergoes a conformational transition from a flexible, square-shaped apo assembly to a compact rhomboid configuration stabilized by extensive inter-subunit contacts (Supplementary Movies 1 and 2). In this active state, the angle between the BC and CT domains narrows by approximately 15°, reorienting the quaternary geometry and reinforcing the domain connectivity. Acetyl-CoA occupies a conserved allosteric site at the BC-PT interface, where it acts as a molecular clamp, stabilizing interdomain hinges, maintaining tetrameric integrity, and preventing BC dimer dissociation, as observed in its absence in both the CT-PT-BCCP complex and apo HsPC^20^. Consistent with this conformational stabilization, acetyl-CoA binding enhances catalytic efficiency by increasing *k*_cat_ from 64.1 to 77.7 s⁻¹ and lowering *K*_m_ from 2.31 to 1.49 mM, corresponding to an overall ∼1.4-fold increase in specific activity. These results confirm that acetyl-CoA binding improves catalytic efficiency not by directly accelerating chemistry but by allosterically stabilizing the catalytically competent conformation and promoting more effective coupling between the BC and CT half-reactions. This supports the flexibility-to-rigidity model of allosteric activation, wherein acetyl-CoA binding reduces the entropic cost of organizing mobile BC domains into a functionally competent assembly^18^. Notably, MtPC exhibits a unique PT-PT interface that forms a reciprocal seam reinforced by hydrogen bonds and salt bridges, in contrast to the weak van der Waals contacts reported in other PC homologs^8^. These observations suggest that in MtPC, the PT domain serves not only as a static scaffold for tetramerization but also as a dynamic hinge modulated by acetyl-CoA to stabilize the catalytically engaged architecture.

Our 3D classification revealed three distinct conformations of the MtPC tetramer, each displaying all four BCCP domains engaged in different arrangements, thereby capturing the possible pathways for BCCP translocation during catalysis. These conformations differ primarily in the positioning of the BCCP domains relative to the two active site pairs (sites 1 and 2) on the opposing faces of the tetramer. In one group of conformations (C8 and C9), the BCCPs were exclusively engaged at either site 1 or site 2 on both faces. Owing to the symmetry of the tetramer, these two conformations are structurally equivalent and indistinguishable by orientation alone. In this state, only one BC dimer, formed at the BC–BC interface, is simultaneously occupied, while the corresponding CT dimer, linked via the CT-CT interface, engages the BCCP on only one face of the tetramer. This arrangement suggests a positively cooperative relationship between the BC dimers, wherein BCCP binding is synchronized at opposing BC active sites, consistent with the observations in LiPC^17^. Conversely, the CT dimers in this state exhibit negative cooperativity; BCCP engagement on one face appears to exclude simultaneous binding on the opposite CT active site, likely coordinating sequential carboxyl transfer and preventing unproductive dual occupancy. In contrast, the third conformation (C4) displayed BCCP engagement at site 1 on one face and site 2 on the opposite face. In this configuration, only one BC domain from each BC dimer is occupied, alternating between the two sites across the tetramer, which is consistent with the half-of-the-sites reactivity model^31^ and reflects negative inter-subunit cooperativity^32^. Notably, this state also reveals a form of positive cooperativity within the CT dimer, wherein both CT domains simultaneously engage BCCPs on opposite faces of the tetramer, suggesting that CT dimers toggle between cooperative modes depending on the overall conformational state.

The coexistence of both positively and negatively cooperative states within a single structural dataset underscores the conformational plasticity of the MtPC. Rather than operating through a fixed mechanistic mode, the enzyme appears capable of dynamically shifting between cooperative regimes, employing positive cooperativity to promote synchronized catalysis and negative cooperativity to enable the alternating engagement of active sites. This regulatory flexibility may reflect an adaptive mechanism that allows MtPC to fine-tune its catalytic output in response to the variable metabolic conditions encountered within the host. However, further studies are required to elucidate the functional basis and physiological significance of these coexisting conformational states.

Sites 1 and 2 are structurally equivalent interfaces, each comprising a pair of BCCP domains arranged head-to-head and docked onto adjacent BC and CT active sites, respectively. Through focused signal subtraction and local refinement, we resolved the high-resolution features of these regions, capturing both catalytic steps of the MtPC reaction: biotin carboxylation at the BC site and carboxyl transfer at the CT site, within a single structural snapshot. This approach also enabled us to trace each BCCP domain back to its originating monomer, revealing that individual BCCPs shuttle exclusively between the two catalytic centers on the same face of the tetramers.

Our findings contrast with the prevailing hypothesis that BCCP engages the CT active site of an adjacent monomer, thereby requiring tetramerization for its catalytic function^8^. However, in MtPC, we observed that BCCP domains interact exclusively with their own BC and CT sites, indicating an intra-monomeric catalytic arrangement. The BC and CT domains in PCs are connected by a single, flexible linker, rendering an isolated monomer structurally unstable for coordinated catalysis. We propose that tetramerization serves to stabilize individual domains and align them into a functionally competent architecture while preserving the interdomain flexibility necessary for BCCP translocation. This structural balance is further modulated by acetyl-CoA, which reinforces inter-subunit contacts and synchronizes the conformational changes essential for efficient turnover.

Focusing on the catalytic site details, in the BC-BCCP engaged state, ADP is well resolved within the BC active site, and biotin, covalently attached to K1093 of BCCP and tethered to the same monomer’s PT domain, adopts an orientation toward R292, forming a hydrogen bond that is absent in *E. coli* BC or LiPC. While this biotin pose typically corresponds to an ATP-bound state in these systems, our structure contains ADP, lacks magnesium density, and shows β-phosphate coordination by E288 and E275. A concurrent salt bridge between R341 and E240 supports the completion of the carboxylation step and provides structural evidence for the proposed gatekeeping mechanism that prevents premature decarboxylation^30^. Although biotin density was weak for explicitly modeling carboxybiotin, the features were consistent with a post-carboxylation intermediate. In the CT–BCCP engaged state, the BCCP domain inserts into the CT active site, with the CT lid closed and biotin positioned near pyruvate and the catalytic Zn²⁺ ion, consistent with catalytically active conformations observed in other PCs^8,17,20,22^. These two snapshots captured sequential catalytic intermediates, demonstrating that MtPC monomers can occupy distinct stages of the reaction cycle simultaneously.

Prior to our study, the positioning of the BCCP domain at the catalytic sites had been inferred from static crystal or cryo-EM structures, which captured it in discrete conformations but failed to resolve the continuous density linking it to the parent monomer due to intrinsic flexibility. Kinetic studies have previously suggested plausible translocation pathways^17^, however, they lack direct structural validation. Using 3DVA, we provide the first direct mechanistic visualization of BCCP movement, revealing how it bridges the distant BC and CT active sites via coordinated conformational changes across the MtPC tetramer. Analysis of the first principal component indicated that pyruvate binding initiates the structural rearrangements necessary for carboxybiotin pocket formation^17,22^. Interestingly, as pyruvate was converted to oxaloacetate, the CT lid tilted inwards towards the tetramer core, coinciding with the loss of detectable pyruvate density (Fig. 6c). This motion likely facilitates oxaloacetate release and propagates an allosteric signal that drives the inward tilting of the opposing BC dimer. These coupled domain movements reposition the BCCP domain at the BC active site, thereby preparing the enzyme for the subsequent catalytic cycle. This observation suggests that BCCP translocation is not a stochastic process but is driven by coordinated ATP and pyruvate binding events that synchronize active site engagement across the tetramer. Complementing these localized transitions, the second principal mode of variability captures the coordinated bending between adjacent monomers that accommodates BCCP translocation between sites 1 and 2, consistent with the ensemble of static conformations observed in our 3D classification. Together, these dynamic reconstructions reveal how substrate binding and domain motions orchestrate the enzyme’s catalytic cycle via long-range allosteric coordination.

Owing to the inherent flexibility of the BCCP domain and its loop to the PT domain, high-resolution 3DVA could not fully resolve the complete translocation trajectory. However, the lower-resolution analysis captured the full dynamic range of BCCP motion across the tetramer as described by two principal components (Supplementary Movie 5). Component one highlights the positively cooperative behavior between BC dimers, arising from coordinated BC–CT domain motions and synchronized BCCP engagement at opposing BC sites. Component two reveals alternating BCCP engagement between the two tetramer faces, providing structural support for the “half of the site” reactivity model. In this configuration, steric and allosteric constraints enforce reciprocal catalysis: one BC dimer adopts a closed BCCP-bound state, while the other remains open, echoing the negative inter-subunit correlation observed in LiPC^17^. These analyses further revealed a continuous density linking each BCCP domain to its parent PT domain, confirming that BCCPs can shuttle between the BC and CT sites of their own monomers. The dual-occupancy state captured in our reconstructions represents a key catalytic intermediate bridging the two half-reactions and offers direct structural evidence for a synchronized alternating mechanism that underlies the efficient catalytic cycle of MtPC.

In conclusion, our integrative structural, kinetics and dynamic analysis revealed MtPC as a finely tuned molecular machine, combining a uniquely stabilizing interdomain architecture with a conserved mode of allosteric activation. By capturing the enzyme with BCCP domains simultaneously engaged at both the BC and CT active sites, a conformation not previously visualized, and mapping the continuous trajectory of domain motions via 3D variability analysis, we provide a comprehensive structural view of the MtPC catalytic cycle. These findings advance our understanding of the mechanistic principles underlying PC function and offer a structural framework for dissecting its dynamic regulation in Mtb and related systems in the future.

## Experimental Procedures

### Protein expression and purification

Full-length MtPC (Uniprot ID I6YEU0) was amplified from the genome of Mtb H37Rv genome using Q5 high-fidelity polymerase (NEB). The amplified PCR product was ligated into the pLATE31 vector (Thermo Scientific) using the ligation-independent cloning method and transformed into Top10 competent cells. The sequence-verified clone was transformed into *E. coli* BL21(DE3) star cells and grown at 37 °C for 3 h, after which protein production was induced using 1 mM isopropyl β-D-1-thiogalactopyranoside for 16 hrs overnight at 18 °C. The cells were harvested by centrifugation and stored at −80 °C until further processing. All purification steps were performed at 4 °C. Cells were harvested, resuspended in 20 mM Tris-HCl (pH 7.5), 300 mM NaCl, 5 mM MgCl₂, and 1 mM DTT (binding buffer), and lysed by sonication. The clarified lysate was loaded onto TALON metal affinity resin, washed with binding buffer containing 10 mM imidazole, and eluted with 200 mM imidazole. The eluted protein was concentrated and further purified by size-exclusion chromatography on a Superdex 200 10/300 GL column equilibrated in binding buffer. Peak fraction containing tetrameric MtPC was analyzed by SDS-PAGE and used for cryo-EM analysis.

### Enzymatic assays

PC activity was measured by coupling oxaloacetate formation to NADH oxidation through malate dehydrogenase, with absorbance monitored at 340 nm. Reactions (200 µL) in a microplate format contained 100 mM Tris-HCl (pH 8.0), 10 mM MgCl₂, 10 mM pyruvate, 1 mM ATP, 5 mM NaHCO₃, 0.2 mM NADH, and 5U malate dehydrogenase (MDH). Assays were performed at 25 °C, in the presence or absence of 0.2 mM acetyl-CoA. Reactions were initiated by adding purified MtPC, and the initial velocities were determined from the linear decrease in absorbance at 340 nm.

### Sample preparation for Cryo-EM

Freshly purified MtPC was diluted to 1.2 mg mL⁻¹ in a buffer containing 20 mM Tris-HCl (pH 7.5), 150 mM NaCl, 5 mM MgCl₂, and 1 mM DTT. For the apo MtPC dataset, the samples were vitrified directly without the addition of substrates or cofactors. For the complex dataset, MtPC at the same concentration was incubated on ice for 30 min in the same buffer supplemented with 1 µM acetyl-CoA, 1 mM pyruvate, and 1 mM ATP to generate a catalytically active state. Immediately before vitrification, 10 mM KHCO₃ was added to the solution. In both cases, 3 µL of the sample was applied to glow-discharged Quantifoil Cu R1.2/1.3 300 mesh grids with a thin continuous carbon layer (10 s, 15 mA). The grids were blotted for 4 s at 4 °C and 100% humidity and plunge-frozen into liquid ethane using a Vitrobot Mark IV (Thermo Fisher Scientific).

### Cryo-EM Data Acquisition and Image Processing

Single-particle cryo-EM data were collected using a Titan Krios G3i transmission electron microscope (Thermo Fisher Scientific) operated at 300 kV and equipped with a BioQuantum K3 direct electron detector (Gatan) in counting mode. Automated data collection was performed in EPU at a nominal magnification of 105,000 ×, corresponding to a physical pixel size of 0.86 Å. Movies were acquired with a total exposure of 50 e⁻/Å², using a defocus range of −0.8 to −2.0 μm. In total, 8,400 movies were collected for the substrate-bound MtPC and 7,500 for the apo MtPC samples, respectively.

All image processing was performed using CryoSPARC version 4.6.2^29^. For the substrate-bound MtPC dataset, 8,400 movies were imported and subjected to patch motion correction and patch CTF estimation. Micrographs with abnormal defocus values, defective images, or poor CTF fits (>5 Å) were excluded using the Curate Exposures tool, yielding 7,820 high-quality micrographs for downstream analyses. Particles were initially picked using Topaz Extract^33^ and extracted in 360-pixel boxes and Fourier-cropped to 90 pixels. Following 2D classification, Ab-initio reconstruction^34^ with five classes was used to discard junk particles, and the most probable class was subjected to Non-uniform refinement^35^. The resulting high-quality map, with the four discarded Ab-initio volumes, along with all the particles of the topaz extract, was used in iterative Heterogeneous refinement, which concurrently increased the particle numbers and mitigated the preferred orientation bias. The final Non-uniform refinement in C1 symmetry produced a 2.95 Å reconstruction. Subsequent Reference-based motion correction and Non-uniform refinement in D2 symmetry yielded a final consensus map at 2.7 Å resolution.

Image processing for the apo-MtPC dataset was performed analogously to that of the substrate-bound complex. The primary difference was the use of the blob picker for particle selection. Ab-initio reconstruction effectively separated intact tetramers from dissociated CT–PT–BCCP populations, both of which were subsequently refined using Non-uniform refinement, yielding the final reconstructions. Local resolution maps were calculated in cryoSPARC using the local resolution tool and subsequently visualized and interpreted in ChimeraX^36^.

### Unsupervised 3D classification, Local refinement, and 3DVA

To address resolution loss due to heterogeneity within each asymmetric unit, particle stacks were symmetry-expanded in D2 symmetry, enabling the analysis of the flexible BCCP domain at the level of individual subunits and their variable orientations. The symmetry-expanded particles were subjected to unsupervised 3D classification in CryoSPARC using a solvent mask derived from non-uniform refinement and low-pass filtered at 12 Å. This approach separated the particles into three distinct conformational classes, effectively resolving the BCCP domains docked at the CT and BC active sites.

For local refinement, a custom mask encompassing the BC–BCCP and CT–BCCP junctions of two adjacent monomers was generated. This mask was then applied to the symmetry-expanded particle set to subtract all other densities, isolating only the BC–BCCP and CT–BCCP-connected regions. Local refinement was performed using the “pose/shift Gaussian prior during alignment” option, with recentring and default parameters applied, while keeping the static mask fixed throughout the refinement.

To analyze the conformational variability in MtPC, 3DVA^28^ was performed in cryoSPARC. A stack of 2,145,612 particles from the symmetry expansion, along with the associated non-uniform volume mask, was used to compute the three components (eigenvectors) of the covariance matrix of the particle distribution. Two separate runs were performed with resolutions of 3 and 12 Å. The model series for each eigenvector was generated using the 3DVA display job with 10 frames per series. Supplemental movies depicting the dynamic trajectories were created using ChimeraX

### Model Building

The initial AlphaFold model of MtPC was fitted individually into the tetrameric cryo-EM density of the substrate-bound MtPC using UCSF ChimeraX. The model was then refined through multiple iterative cycles of real-space refinement in Phenix^37^, applying secondary structure and Ramachandran restraints. For the apo-MtPC map, the substrate-bound MtPC model was flexibly fitted in COOT^38^ using Geman-McClure restraints^39^ and all-atom refinement, followed by real-space refinement. The CT–PT–BCCP model was generated by trimming the complex MtPC model, followed by fitting and refinement, whereas the BCCP–CT and BCCP–BC domains were initially fitted into a locally refined map with subsequent real-space refinement. Additionally, the raw maps were sharpened using EMReady^40^ or LocScale^41^, or DeepEMenhancer^42^ for representation. All molecular graphics, analyses, and movies were created using UCSF ChimeraX.

### Data availability

The data that support this study are available from the authors upon request. CryoEM maps generated in this study have been deposited in the Electron Microscopy Data Bank with IDs: EMD-65383 (apo-MtPC); EMD-63875 (substrate-bound MtPC); EMD-64089 (CT-PT-BCCP); EMD-65188(BC-BCCP/CT-BCCP).

The generated atomic models were deposited in the Protein Data Bank with the following codes: 9VVK(apo-MtPC), 9UBG (complex-MtPC), 9UEQ (CT-PT-BCCP), and 9VMH (BC-BCCP/CT-BCCP).

## Author Contributions

U.D. conceived and supervised this study. A.S. carried out molecular cloning, protein expression, purification, and enzyme kinetics assays. S.S. performed the cryo-EM data acquisition. U.D. conducted cryo-EM data processing, structural analysis, and manuscript preparation. A.S. and U.D. contributed to the data interpretation, discussed the results, and approved the final version of the manuscript.

## Acknowledgements

We thank IIT Delhi, supported by the Department of Science and Technology SATHI facility, Government of India, for providing access to the Thermo Fisher 300 kV cryo-EM for data collection. Computational infrastructure support grant was provided to U.D. by the Indian Council of Medical Research, New Delhi (grant no. ISRM/12(26)/2020), and the All India Institute of Medical Sciences, New Delhi. A.S. acknowledges the Department of Biotechnology, Government of India, for the Junior Research Fellowship. U.D. acknowledges the ANRF (formerly SERB), Government of India, for the SIRE Fellowship that enabled his visit to SLAC, Stanford University, and expresses sincere gratitude to Prof. Wah Chiu for his mentorship and support in advancing cryo-EM expertise.

## Supplementary figure legends

**Supplementary Fig. 1. Protein purification and cryo-EM map-model fitting of apo-MtPC and the CT–PT–BCCP subcomplex. a** Gel-filtration chromatography profile of apo-MtPC showing two peaks corresponding to the MtPC tetramer (∼9.5 mL) and the dissociated CT–PT–BCCP fragment (∼15 mL). Inset, SDS–PAGE profile of MtPC purification. The MtPC band corresponds to ∼120 kDa, while the CT–PT–BCCP fragment migrates at ∼67 kDa. **b** Cryo-EM density map of the CT dimer (deep sky blue and light green) from apo-MtPC with the fitted atomic model. **c** Cryo-EM density map of the BC dimer (deep sky blue and yellow) from apo-MtPC with the fitted atomic model. **d** Cryo-EM density of the BCCP domain (blue) at the CT interface (deep sky blue) in the CT–PT–BCCP subcomplex, showing the BCCP domain fitted into the density. **e** Close-up of the CT–BCCP interface highlighting predominantly hydrophobic interactions.

**Supplementary Fig. 2. Cryo-EM data processing workflow of apo-MtPC. a** A total of 7,500 micrographs were imported into CryoSPARC and processed through patch-motion correction and patch-CTF estimation. Micrographs with CTF fit ≤ 5 Å and suitable ice thickness yielded 6,780 micrographs for particle picking. Blob-picking identified 589,195 particles, cleaned through two rounds of 2D classification to 356,440 high-quality particles. Ab initio reconstruction produced two well-defined classes: apo-MtPC (123,722 particles) and the CT–PT–BCCP subcomplex (110,575 particles). Non-uniform refinement yielded 3.85 Å and 3.20 Å maps, respectively. **b** Representative cryo-EM micrograph of apo-MtPC (scale bar 30 nm). **c** 2D class averages showing apo-MtPC tetramers and CT–PT–BCCP subcomplexes. **d–i** GSFSC plots, angular distributions and local-resolution maps for both structures are shown.

**Supplementary Fig. 3. Cryo-EM data processing workflow of the substrate-bound MtPC. a** A total of 8,400 micrographs were imported into CryoSPARC and processed as described for apo-MtPC. After filtering, 7,820 micrographs were retained for particle picking with Topaz. 334,341 high-quality particles were refined to 3.10 Å, further improved through heterogeneous and D₂-symmetry refinements to 2.70 Å. **b** Representative cryo-EM micrograph (scale bar 30 nm). **c** 2D class averages showing clear secondary structure features. **d–f** GSFSC plots, angular distributions and local-resolution maps for the final reconstruction.

**Supplementary Fig. 4. 3D classification of MtPC. a** Symmetry-expanded particles (536,403) were subjected to D2-symmetry 3D classification into ten classes. Classes C4, C6, C8 and C9 displayed clear BCCP density, while C2 and C5 lacked any. Classes C1, C3 and C7 were discarded. Focused 3D classification around BC–BCCP and CT–BCCP interfaces identified class C4 (461,170 particles) refined locally to 2.90 Å. The right inset shows the local resolution map of the final map. **b–d** Real-space map slices of representative classes showing presence (red) or absence (blue) of BCCP density; only class C4 shows distinct BCCP signal. The horizontal colour scale represents negative (blue) to positive (red) density.

**Supplementary Fig. 5. Cryo-EM density maps and atomic models of BC–BCCP and CT–BCCP active sites. a** Cryo-EM density of the BC active site showing the fitted atomic model. Active-site residues are shown as sticks (deep sky blue), ADP in yellow and biotin covalently attached to BCCP K1093 in blue. **b** Cryo-EM density of the CT active site with fitted atomic model shown as sticks (yellow). The catalytic Zn²⁺ ion (purple) is positioned near its coordination site, with biotin (blue) and pyruvate (cyan) modelled within their respective densities.

**Supplementary Fig. 6. 3DVA of MtPC at 12 Å resolution cutoff**. **a** Two extreme trajectories from 3DVA Component 1 (first frame, deep sky blue; tenth frame, yellow) reveal the BCCP domain positioned exclusively at site 1 on both faces in the initial state, transitioning to site 2 on both faces in the final frame, consistent with a positively cooperative mechanism. The PT–BCCP linkers from corresponding monomers are indicated by superscripts. **b**, Component 2 shows the BCCP located at site 1 on the front face and at site 2 on the back face, consistent with a negatively cooperative relationship between the two dimers. See also Supplementary Movie 5 for the complete conformational transitions.

## Movie legends

**Description of Additional Supplementary Files**

**File Name: Supplementary Movie 1**

Description: Morphed movie illustrating the conformational transition between the substrate-bound and apo forms of MtPC. The movie begins with the substrate-bound MtPC and transitions to the apo state, highlighting large-scale domain rearrangements, particularly the inter-domain PT–PT interactions.

**File Name: Supplementary Movie 2**

Description: Morphed movie illustrating the conformational transition of MtPC highlighting the effect of Acetyl-CoA binding at the BC–PT–CT junction. The movie demonstrates how the presence or absence of Acetyl-CoA alters the relative positioning of the BC domains and modifies the BC–CT inter-domain angle, resulting in a compaction of the overall tetrameric architecture in the substrate-bound state compared with the apo form.

**File Name: Supplementary Movie 3**

Description: 3DVA of Component 1 of MtPC, showing front (left) and side (right) views of the conformational dynamics. The movie reveals a see-saw motion of the BC dimers pivoting along the PT hinge, accompanied by reciprocal movements of the CT domains toward and away from the tetramer centre. The movie frames were generated from the 3D variability display in CryoSPARC, reconstructed as a continuous volume series.

**File Name: Supplementary Movie 4**

Description: 3DVA of Component 2 of MtPC, showing front (left) and back (right) views of the conformational dynamics. The front face reveals bending of the two monomers within the same layer toward each other, accompanied by the appearance of diffuse BCCP density. In contrast, the back face shows that monomers in the corresponding layer do not undergo the same coordinated motion. The overall conformational changes appear to be largely governed by BCCP dynamics.

**File Name: Supplementary Movie 5**

Description: 3DVA of MtPC at a 12 Å resolution cutoff, showing Component 1 (top panel, deep sky blue) and Component 2 (bottom panel, pink), with front (left), back (middle) and side (right) views illustrating the conformational dynamics.

