## Supplementary figures and images for "Cryo-EM structures of Mycobacterium tuberculosis pyruvate carboxylase reveal allosteric activation and domain dynamics during catalysis"

### Supplementary Fig. S1

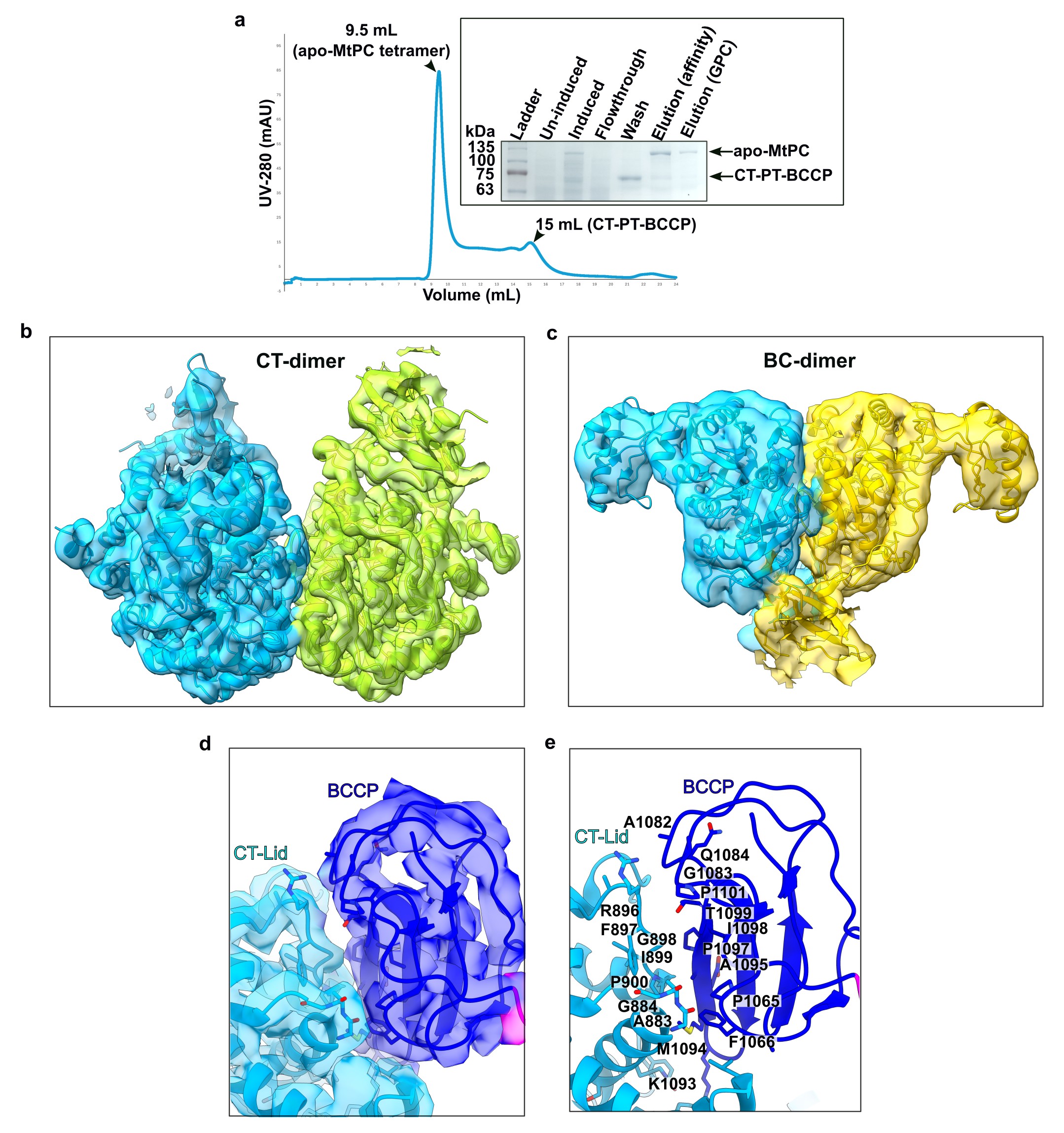

### Supplementary Fig. S2

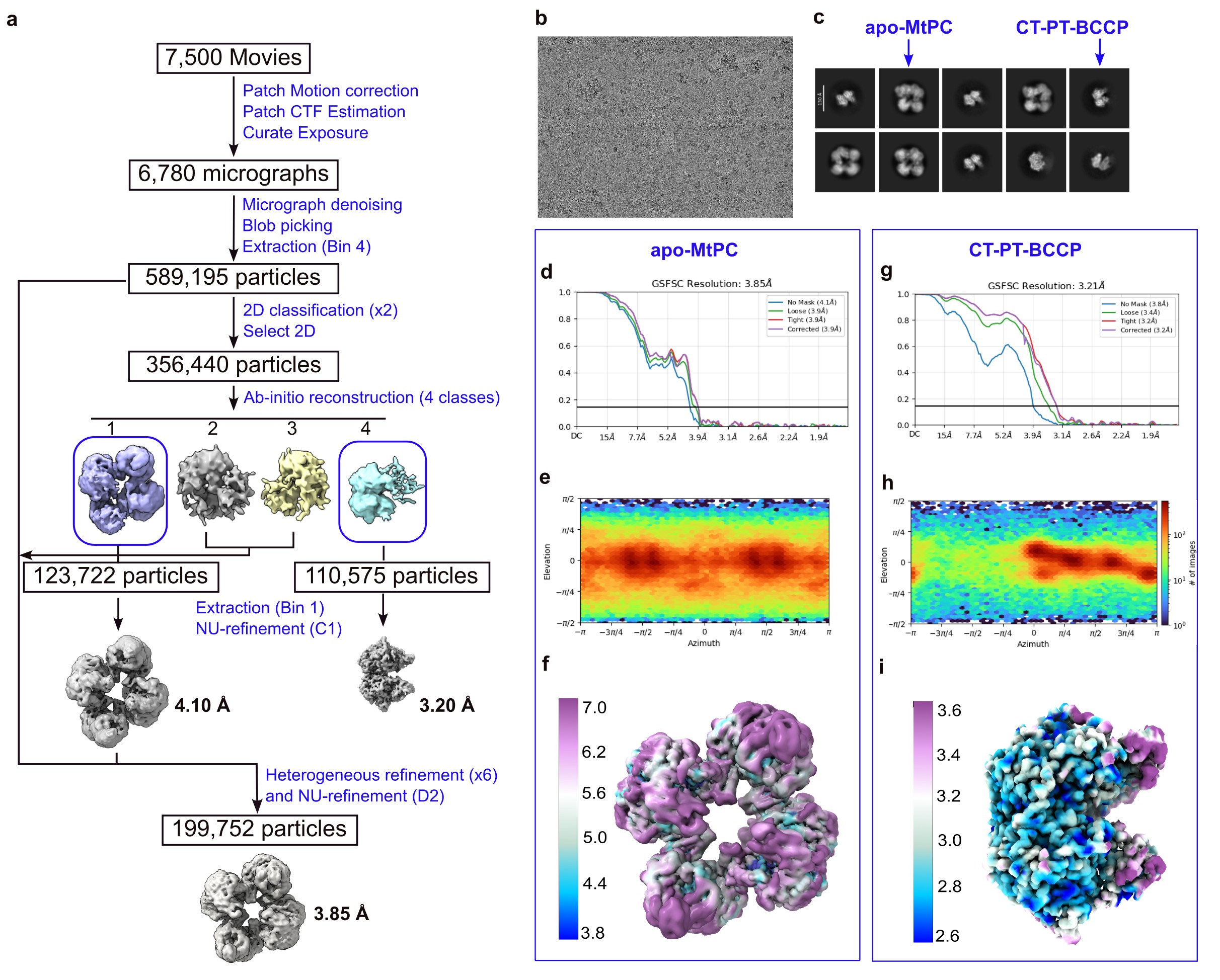

### Supplementary Fig. S3

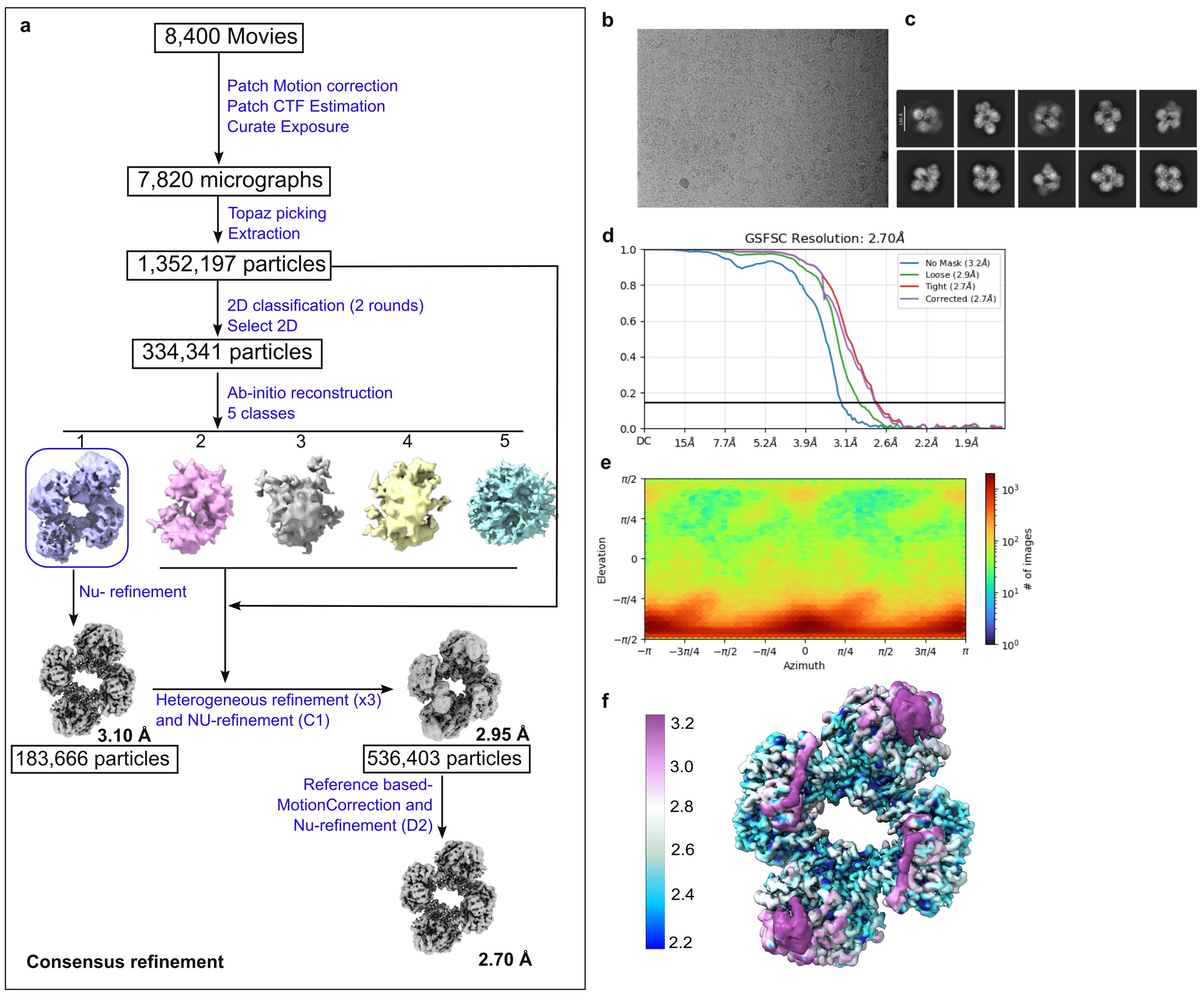

### Supplementary Fig. S4

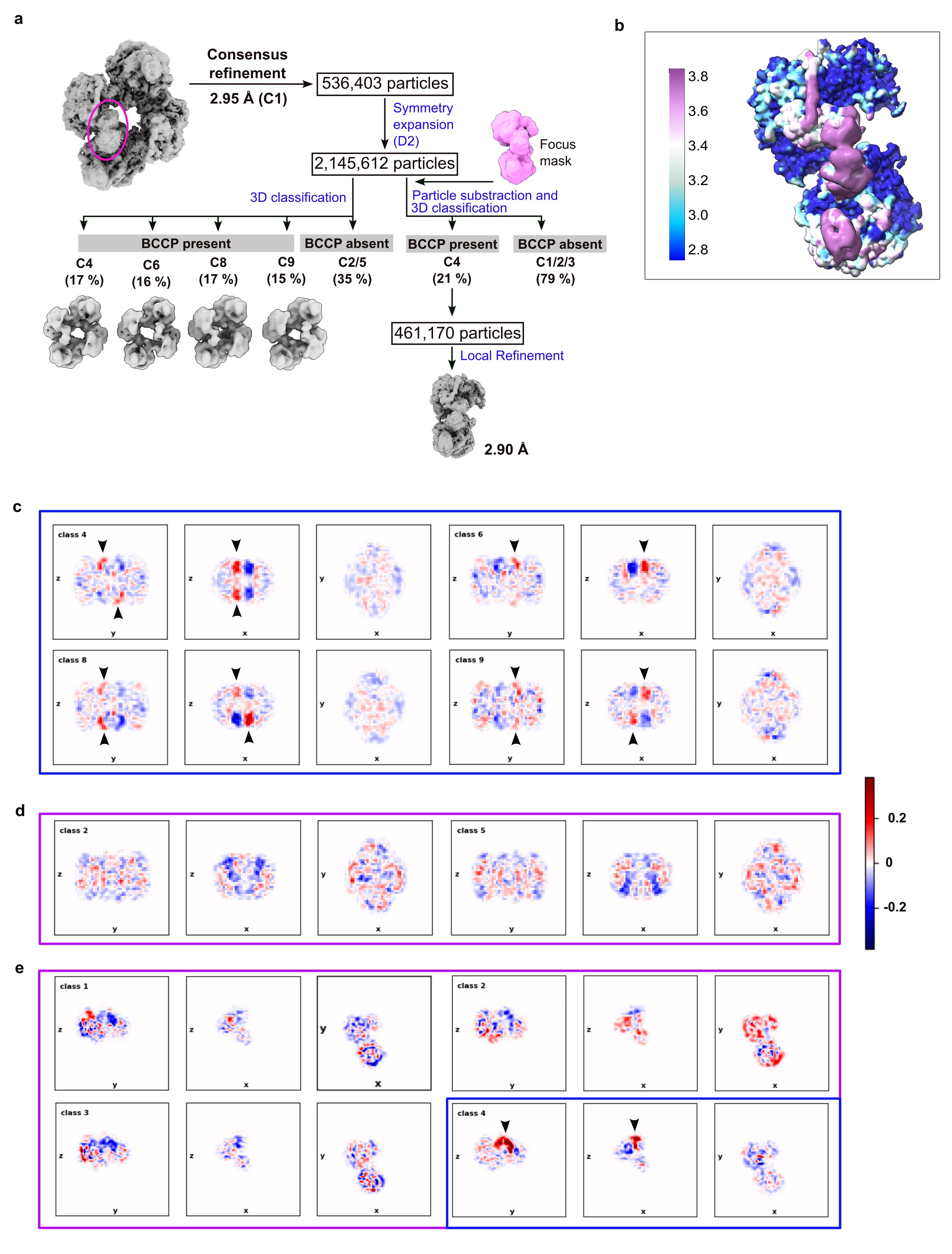

### Supplementary Fig. S5

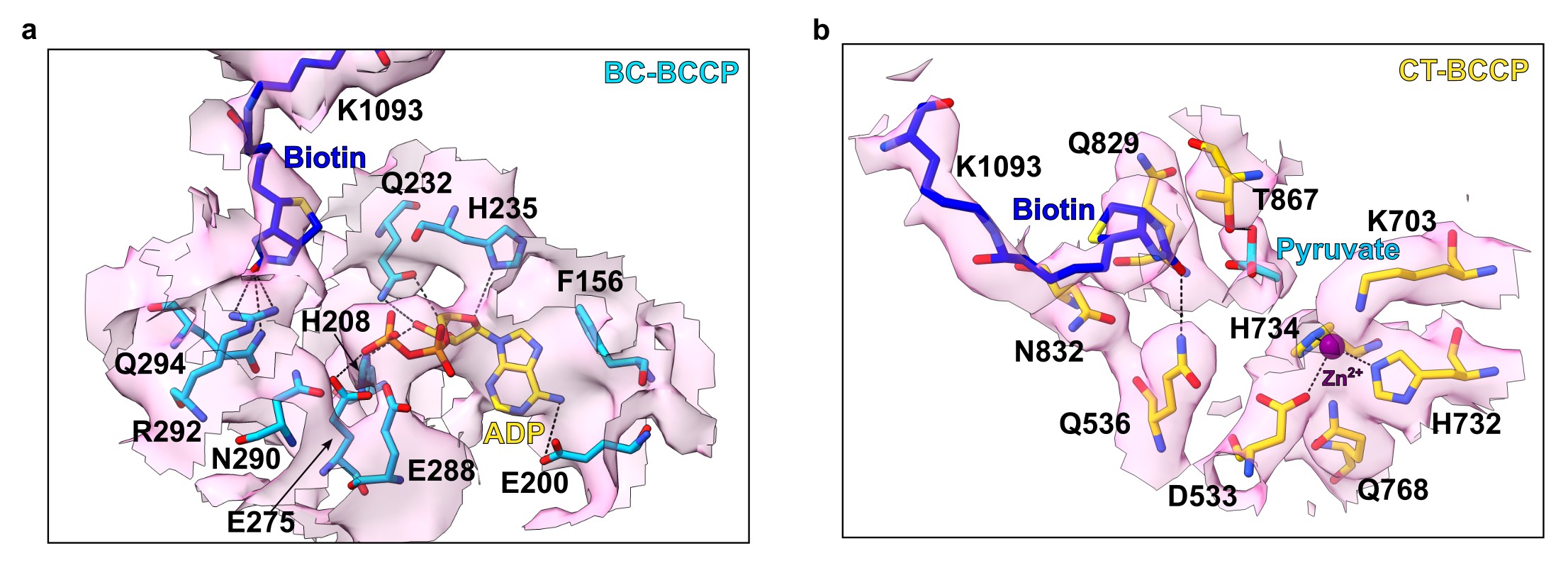

### Supplementary Fig. S6

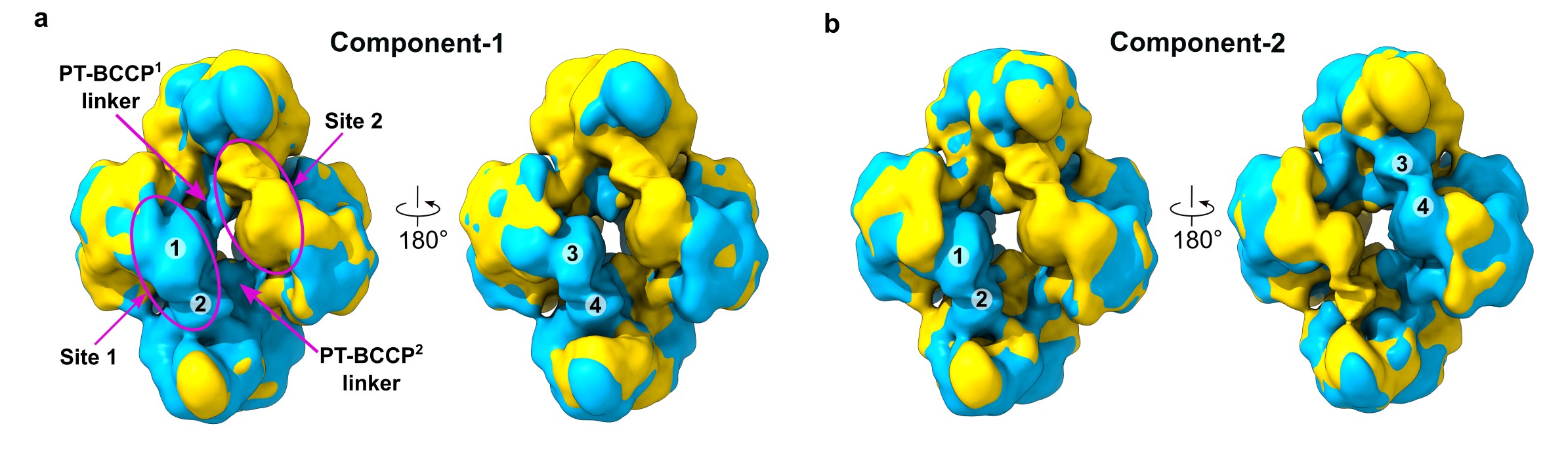
