## Supplementary Table 1 for "Cryo-EM structures of Mycobacterium tuberculosis pyruvate carboxylase reveal allosteric activation and domain dynamics during catalysis"

**Table 1: Data collection and refinement statistics**

|  | #1 Apo-MtPC  (EMDB:65383)  (PDB: 9VVK) | #2: CT-PT-BCCP  (EMDB:64089  (PDB:9UEQ) | #3: Complex MtPC  (EMDB:63875)  (PDB: 9UBG) | #4: BCCP-CT and BCCP-BC  (EMDB:65188)  (PDB: 9VMH) |
| --- | --- | --- | --- | --- |
| **Data collection and processing** | | | | |
| Magnification | 105,000 | 105,000 | 105,000 | 105,000 |
| Voltage (kV) | 300 | 300 | 300 | 300 |
| Electron exposure (e–/Å^2^) | 48 | 48 | 50 | 50 |
| Defocus range (μm) | - 0.8 to - 2.0 | -0.8 to - 2.0 | - 0.8 to - 2.2 | - 0.8 to - 2.2 |
| Pixel size (Å) | 0.86 | 0.86 | 0.86 | 0.86 |
| Symmetry imposed | D2 | C1 | D2 | C1 |
| Initial particle images (no.) | 389,195 | 389,195 | 1,352,197 | 536,403 |
| Final particles images (no.) | 199,725 | 110,575 | 536,403 | 461,170 |
| Map resolution (Å) | 3.8 | 3.2 | 2.7 | 2.9 |
| FSC threshold | 0.143 | 0.143 | 0.143 | 0.143 |
| Map resolution range (Å) | 3.3 - 6.0 | 2.6 - 3.6 | 2.0 - 3.5 | 2.8 - 3.8 |
| **Refinement** | | | | |
| Initial model used | AF | AF | AF | 9UBG |
| Model resolution (Å) | 3.8 | 3.2 | 2.7 | 2.9 |
| FSC threshold | 0.143 | 0.143 | 0.143 | 0.143 |
| Map sharpening B factor (Å^2^) | - 165 | - 55 | - 45 | - 55 |
| Model composition | | | | |
| Non-hydrogen atoms | 31,314 | 19,977 | 33,685 | 137,000 |
| Protein residues | 4,128 | 1,342 | 4,180 | 1,250 |
| Ligand | 0 | 5 (ZN:2, BTN: 2. PYR: 1) | 16 (ADP:4, ACO: 4, ZN: 4) | 5 (ZN:2, BTN: 2, ADP: 1, PYR: 1) |
| B-factors (Å^2^) | | | | |
| Protein | 163.75 | 135.27 | 59.48 | 139.88 |
| Ligand | -- | 131.70 | 55.66 | 175.27 |
| R.m.s. deviations | | | | |
| Bond lengths (Å) | 0.006 | 0.004 | 0.005 | 0.005 |
| Bond angles (°) | 1.215 | 0.996 | 1.169 | 1.014 |
| CC (mask) | 0.68 | 0.82 | 0.82 | 0.82 |
| CC (Volume) | 0.66 | 0.81 | 0.81 | 0.80 |
| Validation | | | | |
| Map-model fit (Q-score) | 0.26 | 0.50 | 0.50 | 0.44 |
| MolProbity score | 1.99 | 1.72 | 1.87 | 2.07 |
| Clash score | 9.69 | 4.10 | 5.04 | 14.34 |
| Rotamers outliers (%) | 0.87 | 0.10 | 1.53 | 0.21 |
| C-beta deviations | 0 | 0 | 0 | 0 |
| Ramachandran plot | | | | |
| Favored (%) | 92.41 | 90.73 | 92.62 | 93.89 |
| Allowed (%) | 7.51 | 9.27 | 7.38 | 6.11 |
| Disallowed (%) | 0.07 | 0.00 | 0.00 | 0.00 |
